# Solar fields in farmlands, their impact on bat presence and activity

**DOI:** 10.1101/2025.10.15.682681

**Authors:** Chloé Tavernier, Rascha Nuijten, Ralph Buij, Frank van Langevelde

## Abstract

Bat populations are facing numerous challenges due to human activities, and the development of solar energy in agricultural landscapes may add to these issues. Effects of solar fields on bat populations are still poorly explored. In this study, we compared bat activity in six solar fields that each had two grassland controls (agricultural grassland and natural meadows). Using passive bat detectors, bat call passes were recorded along the edges of these plots, where the highest bat species diversity and activity were expected. The effects of landscape composition and configuration on the diversity, presence, and activity of different bat species were evaluated for each plot type at yearly, seasonal, and monthly scales. All seven bat species studied were less active in the solar fields compared to the two grassland controls. Specifically, both the number of nights that bats were present and the activity of bats were reduced in solar fields. This study shows that present solar fields have a negative impact on bat activity, which could have population effects as the number of solar fields in the agricultural landscape keeps increasing.

## Introduction

Globally, bats face numerous threats from human activities, with agriculture and energy production ranking respectively as the second and fourth most significant contributors to their decline (1). In particular, insectivorous bat species are negatively affected by intensive agricultural practices, as studies have shown that their foraging activity is lower on conventionally managed farmland compared to organic farmland, likely due to insecticide use (2). In addition to foraging habitat loss, bats are directly impacted by various human activities. For example, in Europe, bat mortality caused by wind turbines is well-documented (3,4). Beyond the effects of wind turbines on local bat populations, there is growing concern that fatalities among migratory bat species could elevate their risk of extinction (1). Bat species are under legal protection in many countries (5,6).

Besides the use of wind turbines, solar energy capacity is growing worldwide (7). However, the impact of photovoltaic fields on biodiversity remains largely unexplored (8,9). Their primary impact on biodiversity is expected to mainly result from habitat modification (10,11). In the Netherlands, for example, where 52% of the land is used for agriculture, solar fields are predominantly established within conventional agricultural landscapes (12), and this trend is the same at the European scale (13). Current studies suggest that solar fields have mainly negative effects on bat populations. In particular, bats exhibit significantly lower foraging activity within solar fields compared to nearby control sites (14). Moreover, it has also been found that solar fields can contribute to bat mortality (15). So far, research comparing bat activity in solar fields to local controls has, however, yielded mixed results. In southwest England, for example, most bat species showed reduced activity in solar fields compared to grassland controls, both within the fields and along adjacent hedgerows (16). In contrast, a study in Hungary found that most bats, with the exception of *Myotis* spp., exhibited similar activity levels in solar fields and grassland controls (17). These differences likely reflect species-specific traits, such as variation in foraging habitats, susceptibility to predation, and flight capabilities as well as local agriculture systems.

Nevertheless, converting intensively farmed land into a solar field could offer potential benefits for bats, as found for other mammal species (Tavernier et al., forthcoming). Indeed, unlike agricultural production, solar fields do not require pesticides or herbicides, which could promote local insect diversity (18). Additionally, most solar fields are enclosed by hedgerows to minimise visual pollution in the surrounding landscape. Hedgerows, often absent in intensively farmed areas, play a crucial role in bat dispersal and habitat connectivity (19,20). The integration of solar fields into homogeneous agricultural landscapes could, therefore, contribute to the restoration of this bat species’ habitat.

Both studies by Tinsley et al. (2023) and Szabadi et al. (2023) were conducted during the summer months (July to October and July to September, respectively). Notably, this period coincides with peak bat activity in agricultural landscapes, as bats have been shown to increase their use of farmland after July (21). However, spring is also a critical season, as bats shift their habitat selection during lactation (22). It is, therefore, possible that solar fields are used at varying intensities across seasons, potentially serving as alternative habitats when agricultural activity fluctuates in surrounding fields. Thus, this study aimed to assess the potential effects of solar fields on bat diversity, yearly bat occurrence, seasonal (spring, summer, autumn) and monthly bat activity. In particular, as bat species may respond differently to the presence of solar fields due to their distinct ecology, the study examines the impact at the species level. To evaluate this impact, occurrence and activity in solar fields were compared with those in two control fields, an intensively managed grassland and a protected meadow. As solar fields are located in farmlands, the two control types were selected as they might represent a possible alternative to land-use from solar fields in the Netherlands.

## Method

### Sampling design

A block-plot design was implemented at six locations to evaluate the impact of solar fields on bat species diversity, presence and activity in the Netherlands. The solar fields were managed by different developers, leading to variations in vegetation management and panel layout. Access was restricted in all solar fields, and they were all surrounded by hedgerows in various stages of development (Supporting information A). In each location, three plots were surveyed: a solar field and two grassland controls (Fig.1). The two controls were an intensively managed grassland for dairy production (intensive control) and an extensively managed meadow for nature conservation (extensive control). The controls were located between two and ten kilometres from the solar fields. These distances were to reduce the effect of solar fields on the bat presence in the controls while staying in roughly the same landscape. Each plot was located in the agricultural landscape, meaning that the matrix surrounding each plot was dominated by farmland. Authorisation to place the detector and visit each plot was granted by their owners.

### Bat survey

In each plot, a stand-alone bat detector (Anabat Swift, www.titley-scientific.com) equipped with an omnidirectional microphone (US-O V3 [10 – 140 kHz], www.titley-scientific.com) was placed from April to November in 2024. The detectors were set up to automatically start recording 30 mins before sunset until 30 mins after sunrise in full spectrum. The batteries and SD cards were changed once a month. The detectors were placed on poles or trees at the edge of the plots at a height of two meters above the ground.

Sound files were analysed in two steps. First, automated identification was performed by Kaleidoscope Pro (23) using the European classifier with a conservative threshold. All *Myotis* and *Plecotus* species were also reclassified under *Myotis* spp. and *Plecotus* spp. respectively, as they are known to be difficult to discriminate based on their echolocation (24). Species or noise that could not be identified by the classifier were manually processed using Anabat Insight (25). However, only *Pipistrellus nathusii, Pipistrellus pipistrellus, Pipistrellus pygmaeus, Nyctalus noctula*, and *Eptesicus serotinus* species, as well as *Myotis spp*. and *Plecotus spp*. were considered in this second classification phase (Table 1). Bat calls were then resampled into events. An event was defined as a cluster of calls from the same species less than three seconds apart.

**Table 1:** Species included in the study and their corresponding preferred foraging habitat, as well as their migratory behaviour. Classification followed Barré et al. (2024) and Heim et al. (2016) classifications. Occurrence refers to the number of nights a species was detected. Ratio Activity refers to the total event from one species divided by the total number of events (165724).

| *Species* | *Foraging habitat* | *Migratory behaviour* | *Occurrence* | *Ratio Activity* |
| --- | --- | --- | --- | --- |
| *Eptesicus serotinus* | Open | Sedentary | 551 | 0.026 |
| *Myotis* spp. | Narrow | Regional or<br>Sedentary | 721 | 0.013 |
| *Nyctalus noctula* | Open | Migratory | 884 | 0.077 |
| *Pipistrellus Nathusii* | Edge | Migratory | 11177 | 0.097 |
| *Pipistrellus pipistrellus* | Edge | Sedentary | 12777 | 0.780 |
| *Pipistrellus pygmaeus* | Edge | Migratory or<br>Sedentary | 121 | 0.001 |
| *Plecotus spp.* | Narrow | Sedentary | 419 | 0.006 |

### Landscape and land use data

The area of different land uses essential for bats in three buffer zones was measured to test the effect of the surrounding landscape composition and configuration on bat community in each of the plots (26,27). The three buffers were set at 500 m, one km and two km from each plot based on the minimum distance between two plots and the effects observed in other studies (19,20). We did not tested landscape effects further than two km as this was the minimum distance between the solar field and the two grassland controls, meaning that beyond this distance we could not distinguish the separate effects of surrounding land uses on the plots anymore. Land uses considered in the landscape were “Forest”, “Shrubs” and “Fresh water” as they are particularly relevant for bat movement and foraging activities. They were extracted and reclassified from the land use map, LGN2023 (Hazeu et al., 2023). It was found that areas covered by tall vegetation and connectivity were particularly relevant to explain bat abundance in farmlands (28). Thus, the landscape composition was defined by the amount of habitat present in the buffer zone, and configuration by the distance between two patches of the same habitat, as well as the minimum distance between each plot and the habitats. The metrics were calculated using the *landscapemetric* package in R (29) or QGIS (30).

### Statistical analysis

All the analyses were conducted using R Statistical Software (31). The analyses were conducted on single species or genus (Table 1).

The effect of landscape composition and configuration on species composition was investigated using a redundancy analysis (RDA). The RDA was performed to test the influence of environmental covariates on species composition in the plots. Environmental metrics, as well as the type of pole the detector was attached to (Table 2), were tested for correlation using a 0.8 threshold and collinearity using a variance inflation factor (VIF) below ten. Step selection was performed on the full and empty model using the *ordistep* function from the *vegan* (32) package, with a limit of permutation of P-values < 0.1 for a term to be added in the final model. A generalised linear model (GLM) using the *glmmTMB* package (33) was used to test the effect of selected landscape metrics on each bat species activity (log-transformed), with the plot set as a random factor and a temporal covariate to account for seasonal change.

**Table 2:**
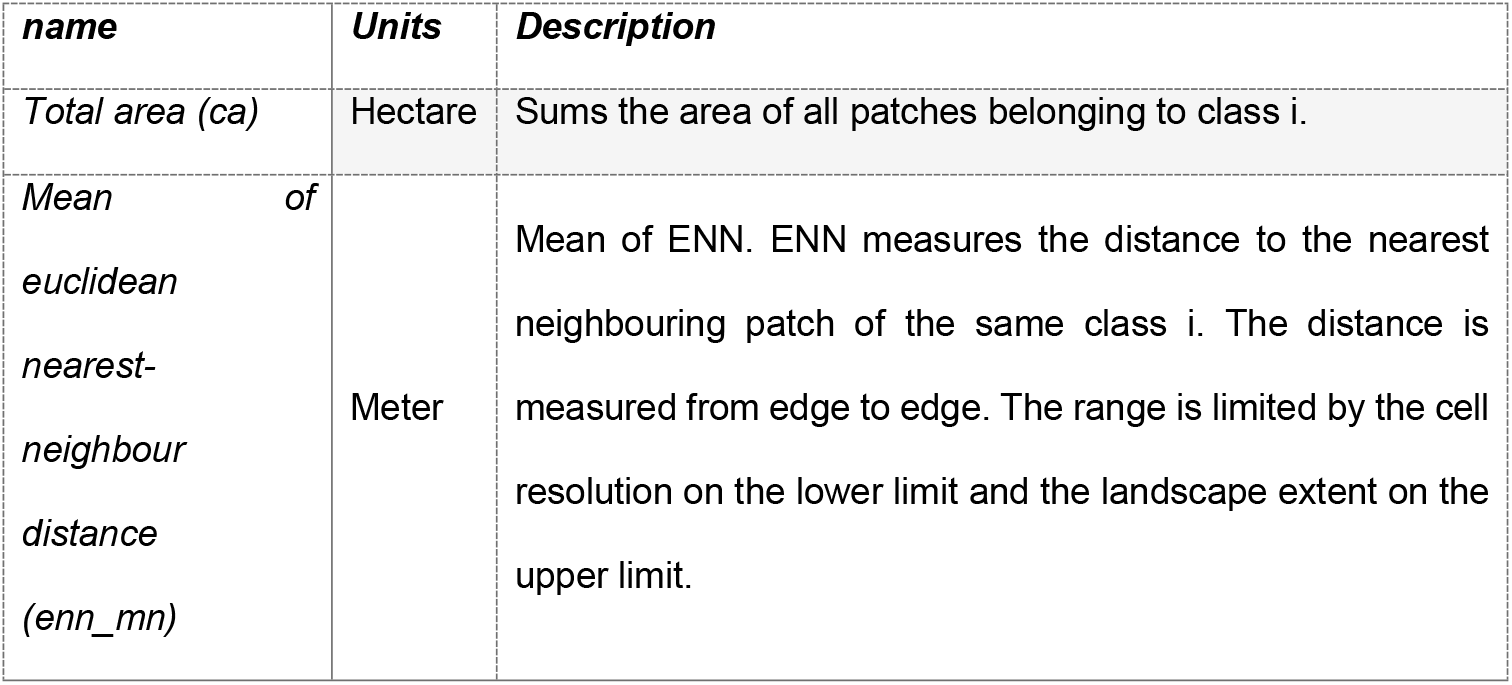
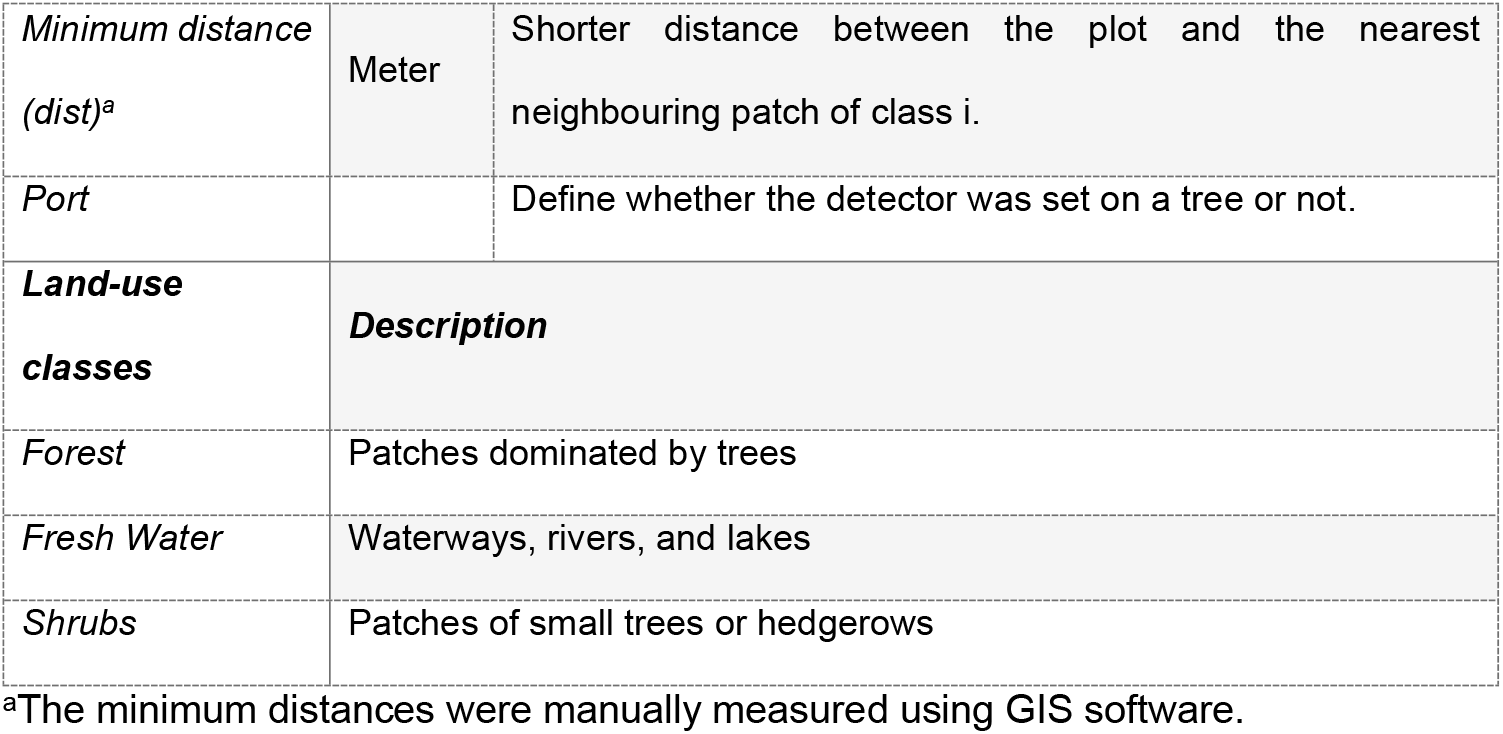
Landscape metrics from the *landscapemetric* package in R (29) and land-use definition used in the present study.

Bat diversity in each plot was measured by Hill numbers for abundance data (equation 1) (34). Three Hill numbers were calculated for each plot with q of order 0 (species richness), 1 (Shannon index), and 2 (Simpson index).

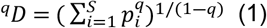

The effect of solar field on bat activity (i.e., number of events per night), bat occurrence (i.e., presence during the night), and bat diversity against the controls was assessed using GLMMs. Bat occurrence and activity were measured at three temporal levels, namely yearly, seasonally, and monthly. GLMMs were fitted with a negative binomial distribution on the activity and with a binomial distribution on the occurrence. Location, the combination of a solar field and its two controls, was included as a random factor. The yearly or monthly efforts (i.e., the logarithm of number of nights the detector was active) were incorporated as an offset, as bat detectors were not active the same number of nights (min = 33 nights, max=153 nights).

## Results

The automatic classifier for bat calls found three species that we did not included in the analyses: *Epstesicus nilssonii, Nyctalus leisleri*, and *Vespertilio murinus*. From the included bats (Table 1), a total of 165 724 events were recorded during the 1318 detector nights (i.e., the number of nights a detector was active) of the survey, with each plot being sampled from 33 to 153 nights between April and November. 78% of the events were records of *P. pipistrellus* (Table 1).

When considering occurrence, no landscape metrics were selected using the RDA during the step-selection that would explain community composition. For activity, the two environmental variables selected for the final RDA to explain community composition were: the distance between forest patches within a two-kilometre buffer area and the distance between freshwater patches within a 500m buffer area. The final RDA, composed of the two selected variables, represented 24.0% of the total variation in species activity between plots. The effect of the distance between forest and water patches was species-specific (Fig.2).

The solar field and the two controls had the same species richness (7 species), meaning that all species visited the three plot types. However, there was strong evidence of a more uneven coverage of species in solar fields due to hyperabundance of *P. pipistrellus*, as indicated by the lower Shannon and Simpson diversity indices compared to the extensive and intensive controls (Fig. 3).

**Fig. 1:**
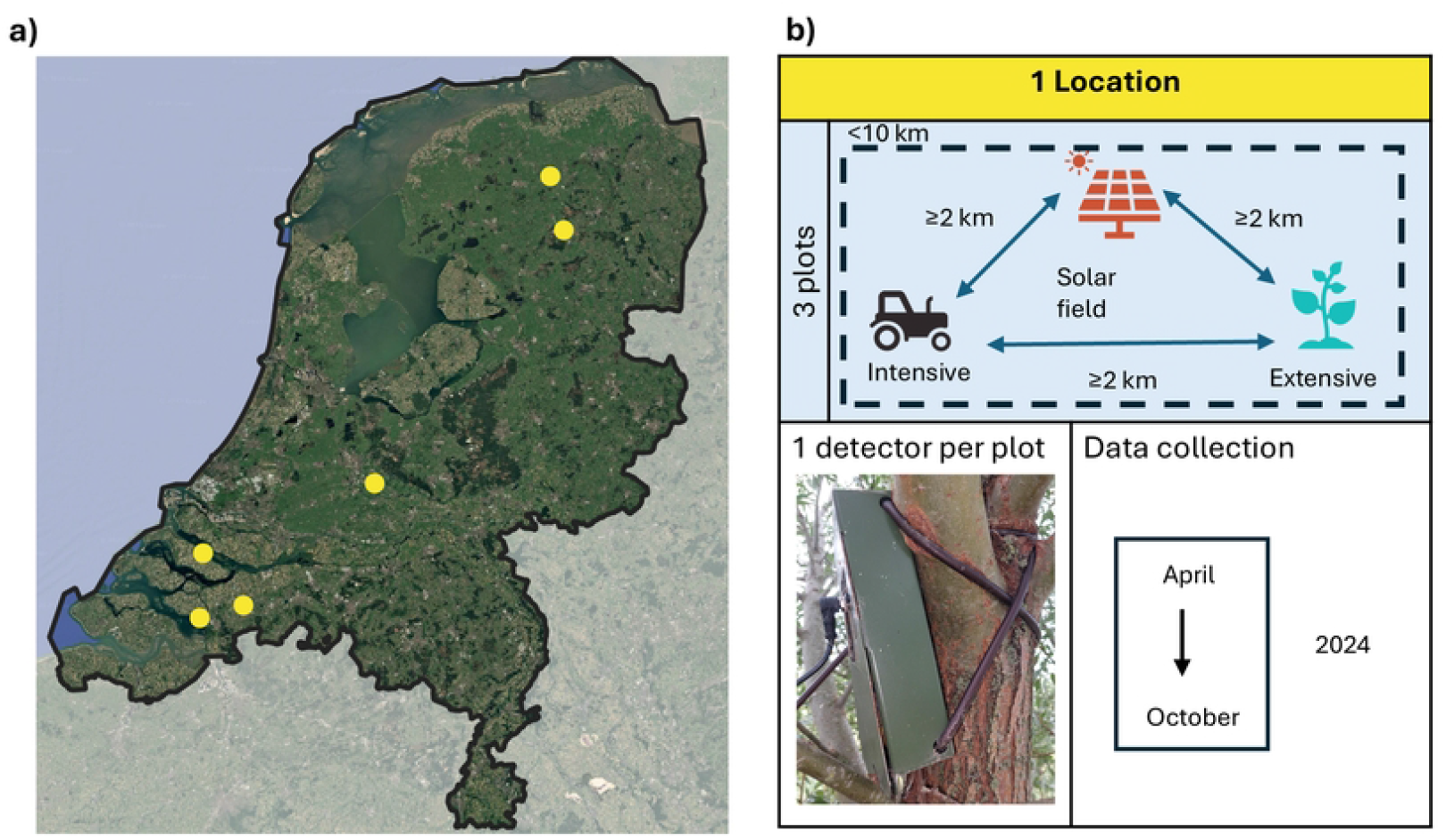
Sample design in seven solar fields and their controls. **(a)** Satellite image of the Netherlands and the six locations where the bat survey took place. **(b)** Description of the survey design, where each location was composed of three plots: the solar field and the two control fields, being an extensive meadow and an intensive grassland.

**Fig. 2:**
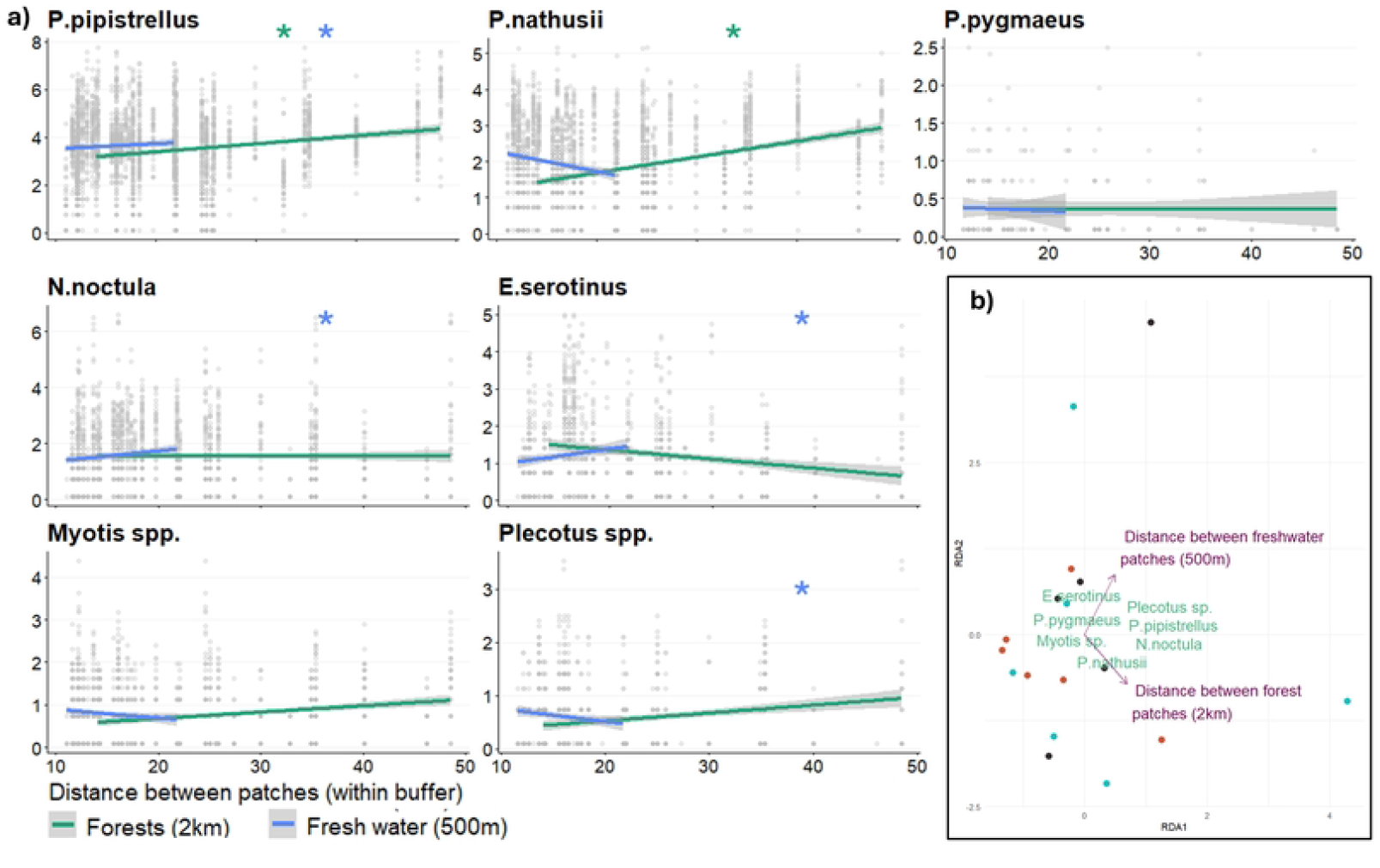
Bats’ activity and the landscape metrics. **a)** Bat activity per night related to the distance between the forest (green) or freshwater patches (blue) in the landscape. Forest patches are within a 2km buffer, while freshwater patches are within a 500m buffer around each sampling plot. Activity is log-transformed, and distance is expressed in meters. Lines show the linear regression. Significance of the landscape metric on activity is expressed by a star. **b)** RDA output of the species composition in different plots. Species are in green in the following format: Genus.species. The constrained metrics explaining 24% of the variation found in species composition are shown in purple. The landscape metrics are calculated in a buffer around each plot; the size of the buffer is in brackets (500m or 2 km).

**Fig. 3:**
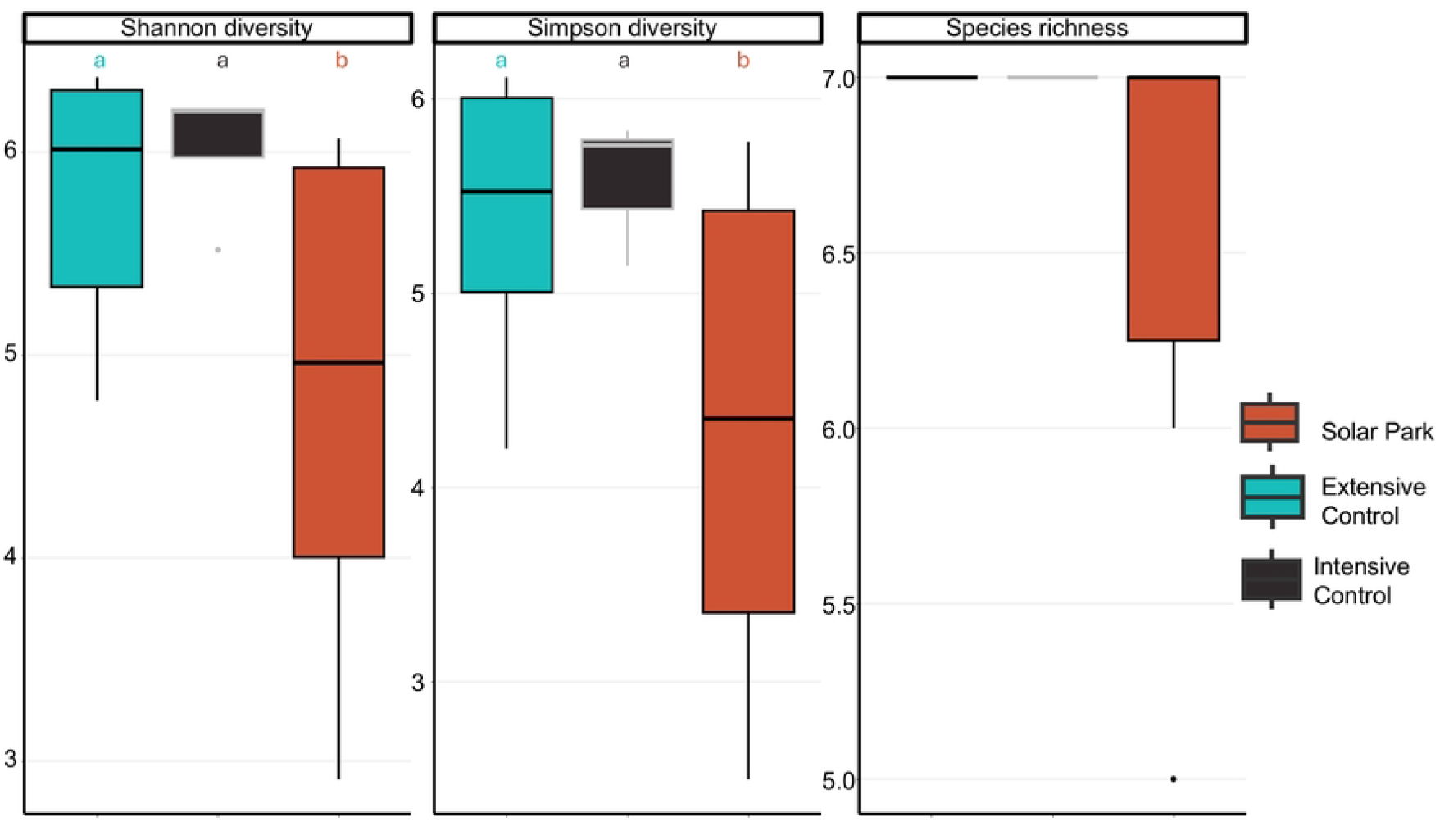
Box plot of the three Hills index. Hill numbers with q = 0 (Species richness), q = 1 (Shannon diversity), and q = 2 (Simpson diversity) calculated in solar fields (in red), extensive controls (in blue), and intensive controls (in black). Significant differences between plot types are indicated by different letters (p < 0.05), with letter order representing the direction of the effect (a > b). No letter indicates no significant difference There was strong evidence of lower nocturnal activity in solar fields than in the two controls for all species. The same pattern was found for their nocturnal occurrence, except for *P*.*pipistrellus* and *Plecotus spp*., where no evidence was found of a difference between plot types (Fig.4).

The two months that were the most sampled were May and June. Solar fields showed a consistently lower activity and occurrence than the two controls at all temporal scales measured than the two controls (Fig. 5).

## Discussion

This study builds upon the growing body of research exploring the impact of solar fields on biodiversity. Specifically, it examines how existing solar fields affect bat activity, species known for their sensitivity to land-use changes and widely recognised as bioindicators of ecosystem health (35). We compared bat diversity, occurrence, and activity in solar fields with those in intensively managed grasslands and extensively managed meadows, habitats known to negatively and positively influence bat populations, respectively (2,19).

In this study, all bat species exhibited reduced presence and activity in solar fields compared to the controls (Figs. 4 and 5). Notably, even intensively managed grasslands demonstrated higher bat activity and presence than solar fields (Fig. 4). Throughout the year, bat activity was highest in extensive meadows and lowest in solar fields, a pattern consistent across all species except for *P. pygmaeus* (Fig. 5). These findings suggest that solar fields in our study are failing to provide potential useful habitat for all bat species.

**Fig. 4:**
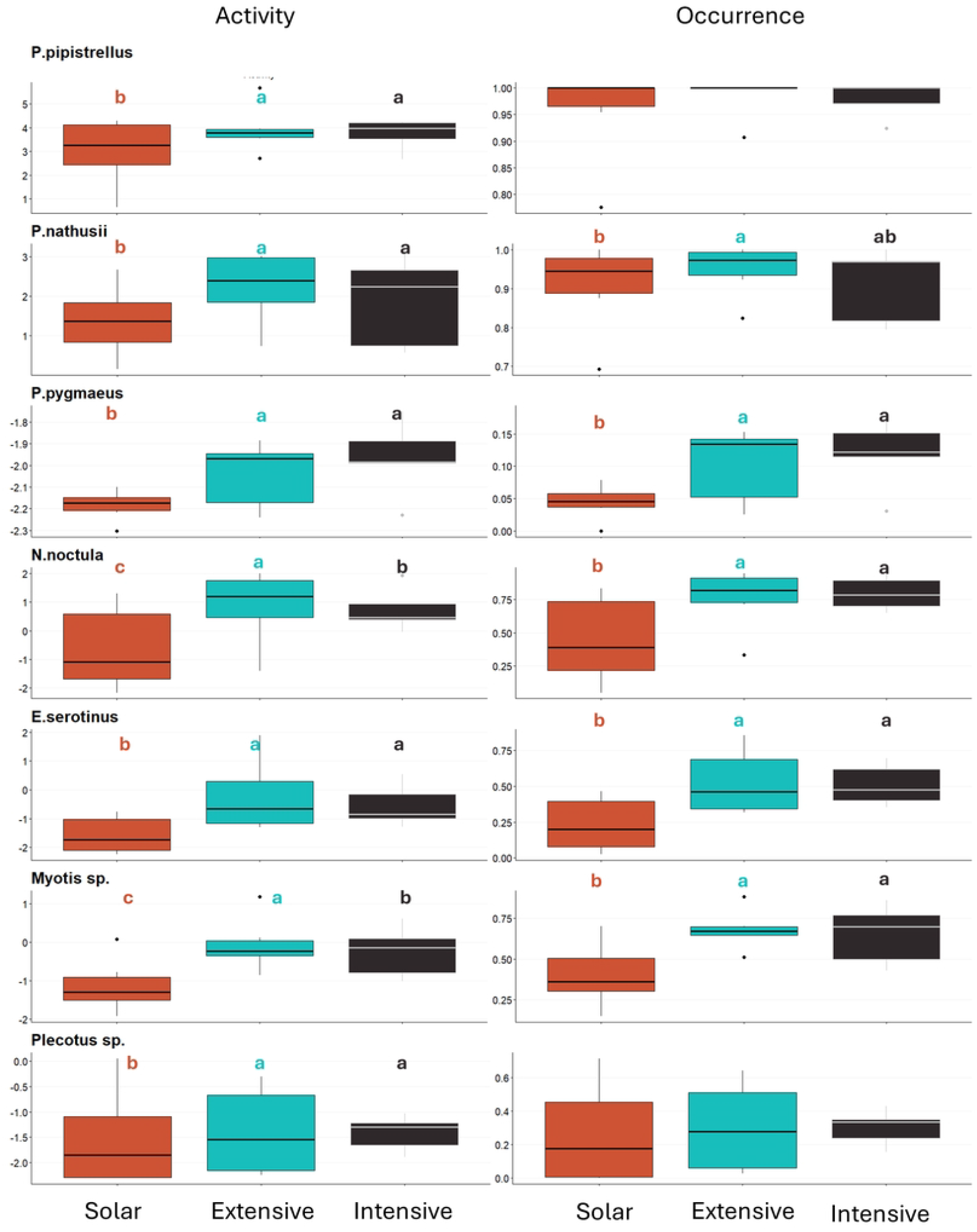
Box plot of activity (left) and occurrence (right) of each species in the three plot types. The plot types are solar fields (in red), extensive controls (in blue), and intensive controls (in black). Significant differences between plot types are indicated by different letters (p < 0.05), with letter order representing the direction of the effect (a > b). No letter indicates no significant difference. Activity (y-axis of the left panels) is log-transformed.

**Fig. 5:**
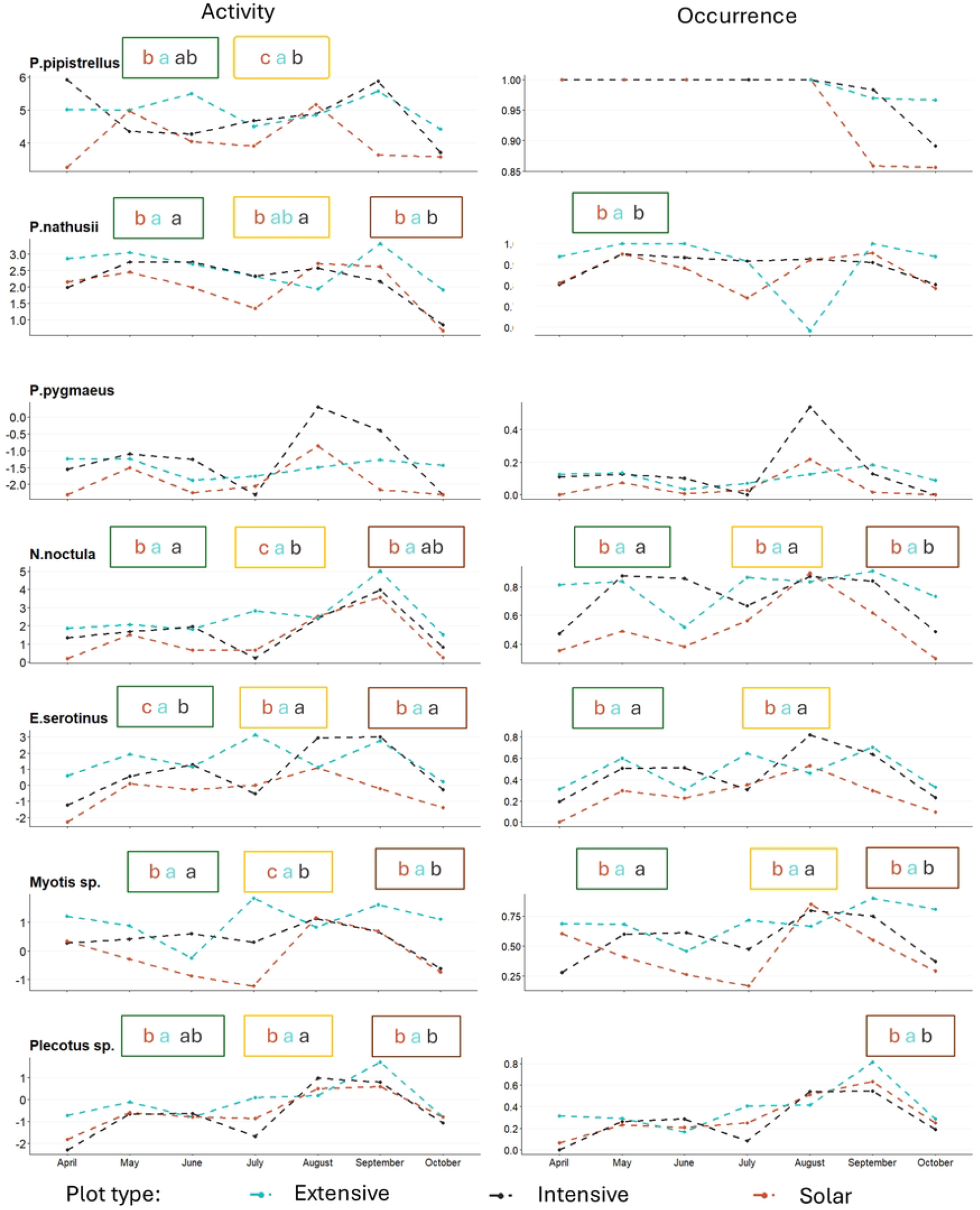
Mean of activity (left) and of occurrence (right) per month of each species in the three plot types. The three plot types are solar fields (in red), extensive controls (in blue), and intensive controls (in black). Significant differences between plot types per season (Spring (April-June) = green, Summer (June to September)= yellow, Autumn (September to October)= brown) are indicated by different letters (p < 0.05), with letter order representing the direction of the effect (a > b). No letter indicates no significant difference. Activity (y-axis of the left panels) is log-transformed.

Two species groups showed a weaker habitat avoidance of solar fields compared with the controls: *P*.*pipistrellus* and *Plecotus spp*.. *P. pipistrellus* is a ubiquitous and highly synanthropic species, and it is the most common bat species in western Europe (36,37). This characteristic might explain its presence in all three plot types on nearly every night surveyed. In contrast, *Plecotus spp*. are strongly associated with forested habitats such as forests, orchards, or parks (36). The focus of this study on grasslands as an alternative habitat for solar fields may explain the limited use of these habitats by *Plecotus spp*. and consequently the absence of occurrence differences between plot types.

### Habitat connectivity is more important than habitat quantity

In this study, the community compositions of bats across the three plot types exhibited considerable homogeneity, as all bat species were present in the three plot types. The study locations were situated within a farmed landscape, characterised by its homogeneity and its scarce amount of natural habitats.

These findings contrast with previous research in the literature. The amount of forest and freshwater patches are well-established predictors of bat diversity and activity (38). However, the specific effects of landscape composition and configuration parameters can be species-specific, complicating the selection of unique landscape metrics to explain the entire bat community composition (Fig.2). For instance, Fuentes-Montemayor et al. (2011) found that *P. pipistrellus* and *P. pygmaeus* exhibited different responses to landscape connectivity and the scales at which parameters were measured. Additionally, the seven bat species of the present study have varied ecologies, such as *N. noctula*, which can travel up to 26 km from their roosts, and *Plecotus* species, which typically remain within five kilometres of their roost. These examples underscore the species-specific nature of landscape effects and the challenges in selecting universal landscape metrics to explain bat community composition. Nevertheless, this study emphasises the need for connectivity between forested and freshwater patches in the landscape.

### Negative impact of solar fields

The present study demonstrates a clear reduction in the occurrence and activity of bats within solar fields in the agricultural landscape of the Netherlands. The findings indicate that solar fields fail to contribute to bat conservation; they even reduce the habitat quality of bats in farmlands. These results are consistent with a previous study comparing the edges of solar fields with the edges of adjacent control sites in the UK (Tinsley et al., 2023). Unless for *Plecotus spp*., Tinsley et al. (2023) also found a reduction in activity at the edges of solar fields compared to the edges of controls for the same bat species as in this study. Potential reasons for the reduction in bat activity cited included a lack of prey (insects) and the collision risk posed by the clutter of solar panels (39).

The hypothesis concerning the lack of prey requires further exploration. Barré et al. (2024) found that bats have a straighter and faster flight behaviour in solar fields associated with a reduction of feeding behaviour. A study measuring insect abundance in the same solar fields as the present study found that solar fields had fewer ground-emerging Diptera and Coleoptera compared to control sites (Kocsis et al., forthcoming). These two orders are known to be important food resources for bats (40). Therefore, a better comprehension of prey availability as well as the bat capabilities to detect and catch them within solar fields is essential to fully understand the impact of solar fields on bat activity. Therefore, to reduce the negative impact of solar fields on bats, it is necessary to enhance their suitability for flying insects while minimising the risk of bat collisions.

### Perspective of solar fields for bat conservation

This study demonstrates that current solar fields do not support the occurrence or activity of bat species. Although developers often commit to implementing mitigation or enhancement measures during construction, these are rarely applied effectively in practice (41). Nevertheless, the negative impacts of solar fields on bats could be reduced through targeted ecological enhancement and by prioritising installations on intensively used farmland. For instance, integrating features such as hedgerows, tree lines, and flower-rich grasslands into solar developments could improve landscape connectivity and boost local insect abundance and diversity (19,20). Solar fields have the potential to address both the energy and biodiversity crises, but at present, they fail to support bat communities, even within the already degraded agricultural environments.

## Acknowledgments

We dearly thank the students (Elske Wijshake, Joep van Dongen, Oscar Creus Fabregat, Thomas Huijskens) who have helped with the annotation of the audio events, as well as Aliki Marmara, Dimitri Tavernier, and Hanna Willers. We express our gratitude to all asset managers, farmers, and Natuurmonumenten and Staatsbosbeheer field managers for facilitating our entry to their fields. Finally, we thank Sebastiaan Forouzan Fard of Eelerwoude for facilitating communication with the solar field managers. Artificial intelligence-generated content was used to improve the clarity and conciseness of the text.

## Statements & Declarations

### Data Availability Statement

All occurrence data are available in Zenodo (42)

### Authors’ contributions

Chloé Tavernier, Rascha Nuijten, Ralph Buij, and Frank van Langevelde conceived the ideas and designed the methodology; Chloé Tavernier lead the collection of the data and analysed them; Chloé Tavernier led the writing of the manuscript. All authors critically contributed to the drafts and gave final approval for publication.

